## Supplementary material for "Hypoxia-enhanced Blood-Brain Barrier Chip recapitulates human barrier function, drug penetration, and antibody shuttling properties"

[1]Wyss Institute for Biologically Inspired Engineering at Harvard University, Boston, MA 02115, USA; [2]Department of Neurosurgery, Boston Children's Hospital and Harvard Medical School, Boston, MA 02115, USA; [3]Department of Chemical and Biological Engineering, University of Wisconsin-Madison; [4]Harvard John A. Paulson School of Engineering and Applied Sciences, Harvard University, Cambridge, MA 02138, USA; and [5]Vascular Biology Program and Department of Surgery, Boston Children's Hospital and Harvard Medical School, Boston, MA 02115, USA.

[#]Co-first authors

[†]Current Address: Ulsan National Institute of Science and Technology (UNIST), UNIST-gil 50, Ulsan 689-798, Republic of Korea

[‡]Current Address: Department of Micro and Nanosystems, KTH Royal Institute of Technology, Stockholm, Sweden and Department of Neuroscience, Karolinska Institute, Stockholm, Sweden

*Corresponding Author: Donald E. Ingber, M.D., Ph.D., Wyss Institute at Harvard University, CLSB5, 3 Blackfan Circle, Boston, MA 02115 (Em:; Ph: 617-432-7044; Fx: 617-432-7828)

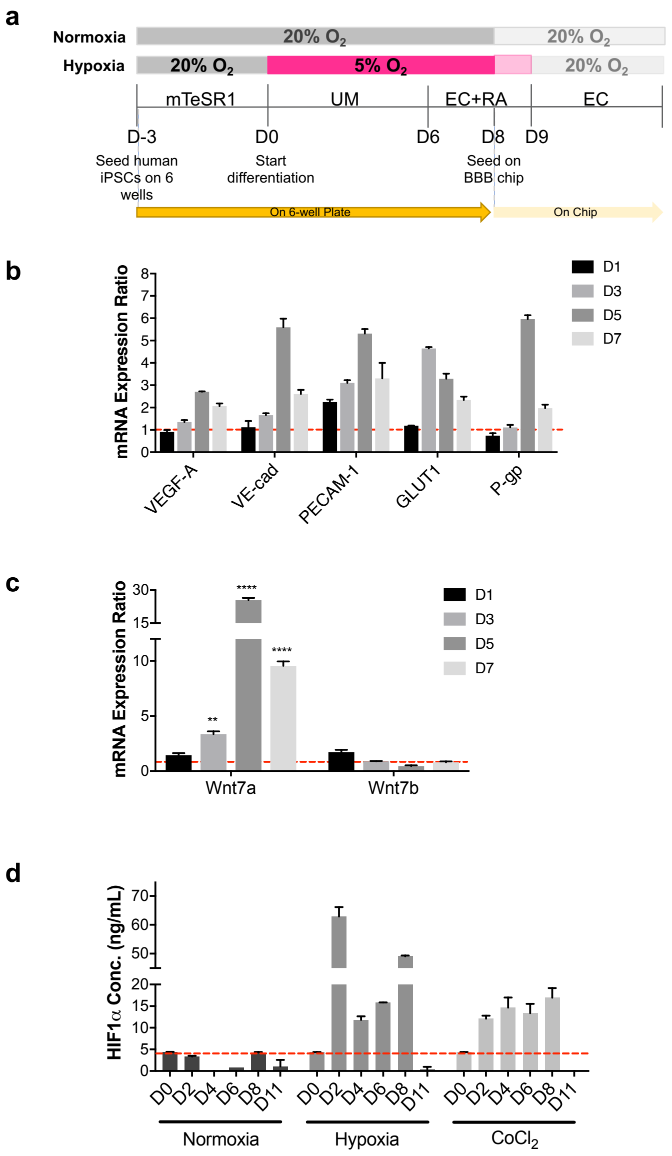


### Supplementary Fig. S1

**(a)** Timeline for the differentiation of the iPS cells to the human BMVECs, and seeding on the BBB chips. **(b)** Fold changes of mRNA expressions of VEGF-A, VE-cadherin, PECAM-1, GLUT1, and P-gp during differentiation (D1-D7) of iPS-hBMVECs under hypoxia relative to normoxia analyzed by qRT-PCR. **(c)** Relative fold change in mRNA expression of Wnt7a and Wnt7b in iPSC differentiated under hypoxic condition compared to normoxia during the differentiation process (D1-D7). **(d)** ELISA analysis for

HIF1$\alpha$ protein expression during differentiation of iPS-BMVEC (D0-D8) and after differentiation (D11).

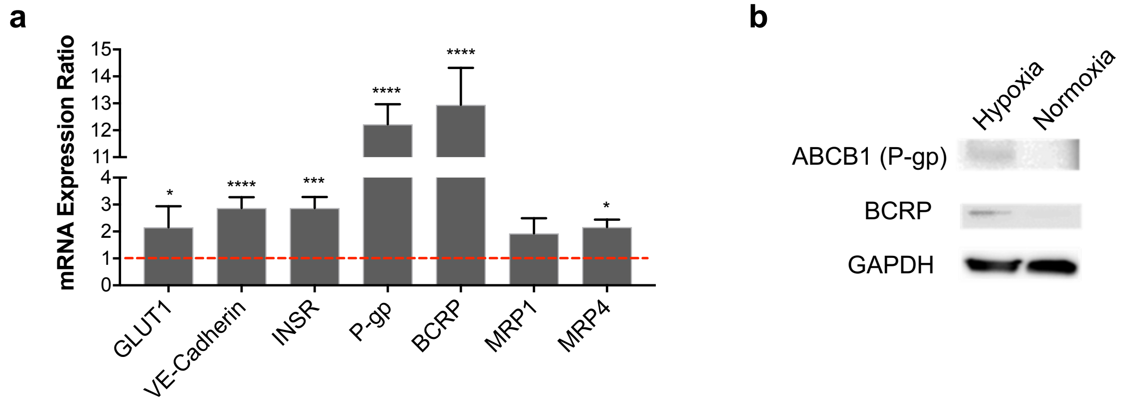


**Supplementary Fig. S2**

**(a)** Relative fold changes in mRNA expressions of GLUT1, VE-cadherin, insulin receptor (INSR), P-gp, BCRP, MRP1, and MRP4 in iPS-BMVEC differentiated under hypoxic versus normoxic conditions. **(b)** P-gp and BCRP protein expression of the iPS-BMVEC differentiated under hypoxia and normoxia were compared using a Western Blot. GAPDH was used as a control protein.

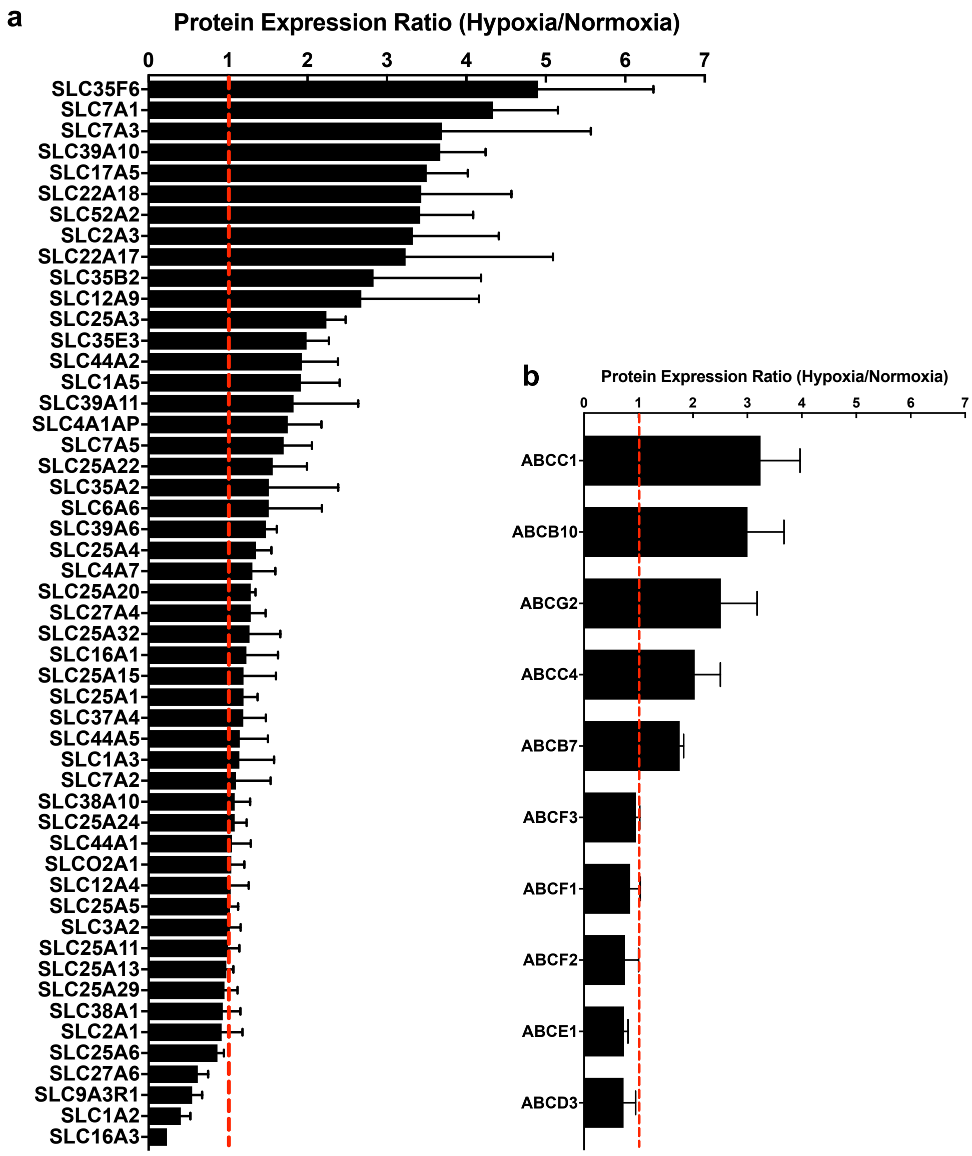


### Supplementary Fig. S3

All SLC proteins **(a)** and all ABC proteins **(b)** identified in the proteomics studies on the iPS-BMVEC differentiated under hypoxia and normoxia. Graphics show the relative abundance of the SLC and ABC proteins in the hBMVECs induced hypoxia vs normoxia conditions.

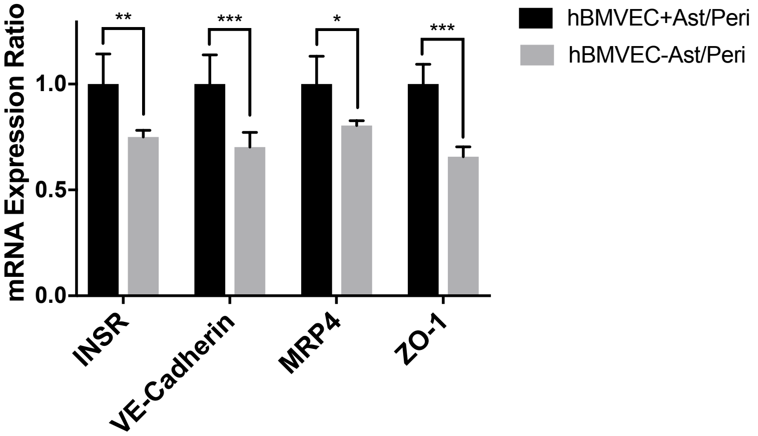


**Supplementary Fig. S4**

mRNA expressions of INSR (insulin receptor protein), VE-cadherin, MRP4, and ZO-1 on the BBB Chips in the presence and absence of astrocyte and pericyte coculture were quantified using qPCR.

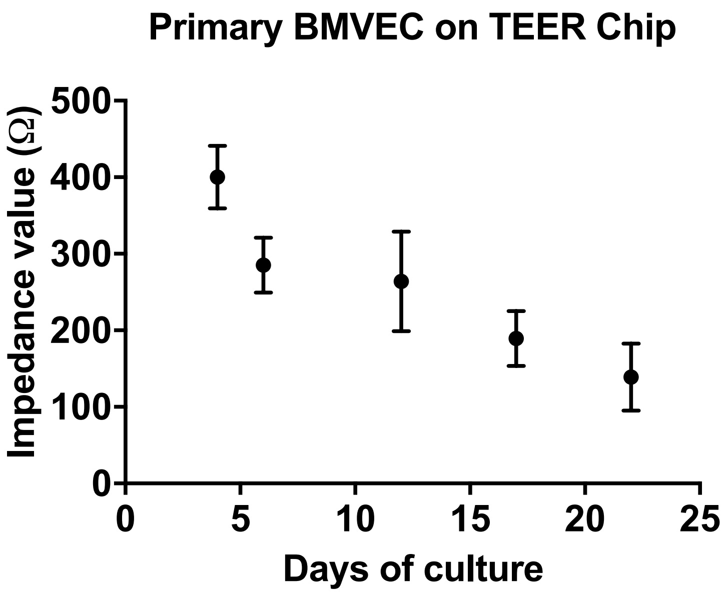


**Supplementary Fig. S5**

Barrier integrity of the primary human BBB Chip monitored in TEER chips with impedance measurements, recorded in the frequency range of 0.1 Hz to 100 kHz over 3 weeks after seeding primary BMVECs along with astrocytes and pericytes.

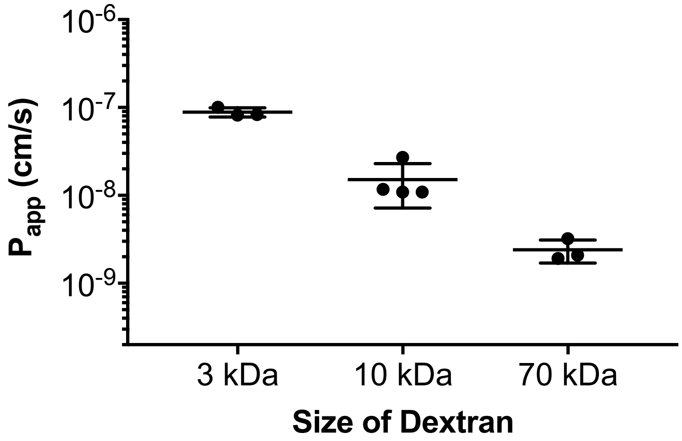


**Supplementary Fig. S6**

Permeability of dextran tracers of various sizes in the BBB Chips. Fluorophore-labelled dextran molecules (3, 10, or 70 kDa) were flowed through the brain channel on the BBB Chips for 3 h at 100 µL/h flow rate. Effluent samples from both brain and vascular channels were collected and fluorescent intensity of the samples were detected, and dextran concentrations were quantified based on standard curves to calculate $P_{app}$ values.

a

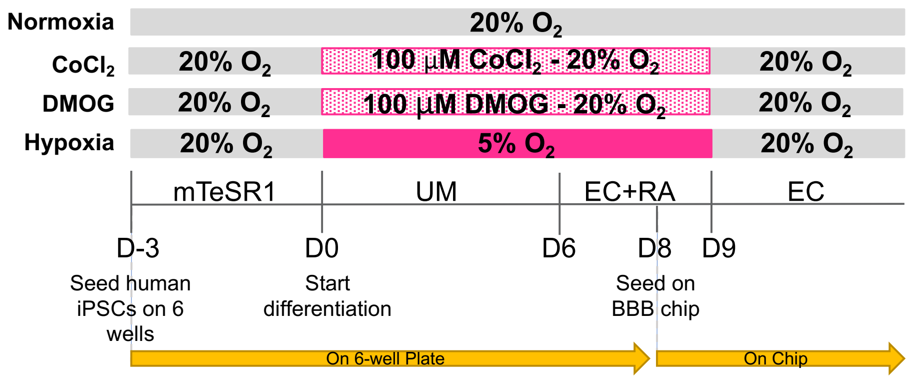


b

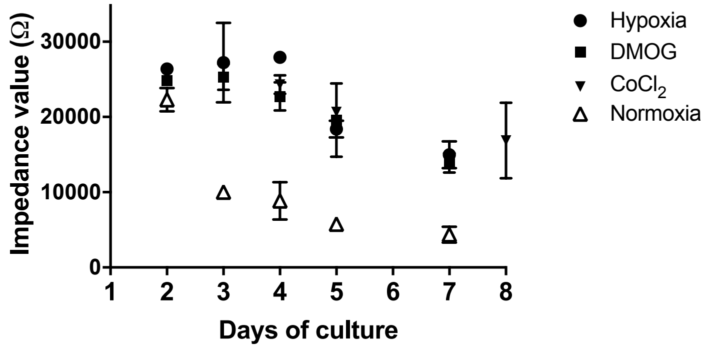


**Supplementary Fig. S7**

**(a)** Experimental design schematic showing how BBB Chips were generated using iPS-BMVECs differentiated under normoxia (as control), hypoxia, or using chemical inducers ($CoCl_2$ and DMOG) that mimic hypoxia under normoxic conditions. **(b)** Impedance measurements of barrier integrity of BBB Chips generated with iPS-BMVECs differentiated as described in **a** and measured in TEER chips.

a

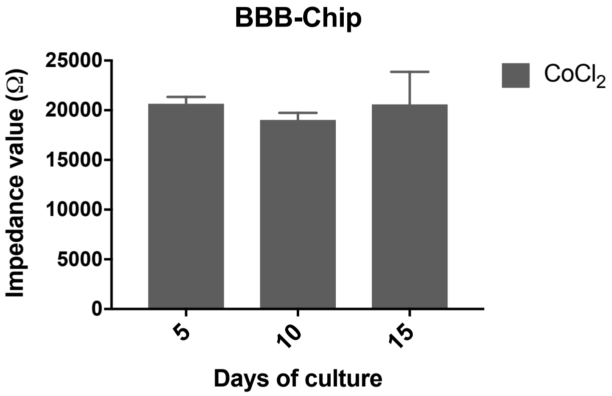


b

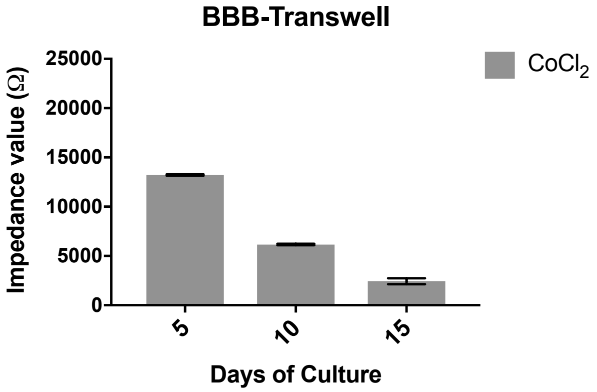


**Supplementary Fig. S8**

The high barrier function (impedance measured by TEER) was sustained for longer times in BBB Chips with iPS-BMVECs that were induced to differentiate using $CoCl_2$ interfaced with astrocytes and pericytes(**a**) and this effect appeared to require flow as it was not observed when the same cells were interfaced in static Transwell cultures **(b).** Barrier integrity on the BBB Transwells was monitored by TEER measurements.

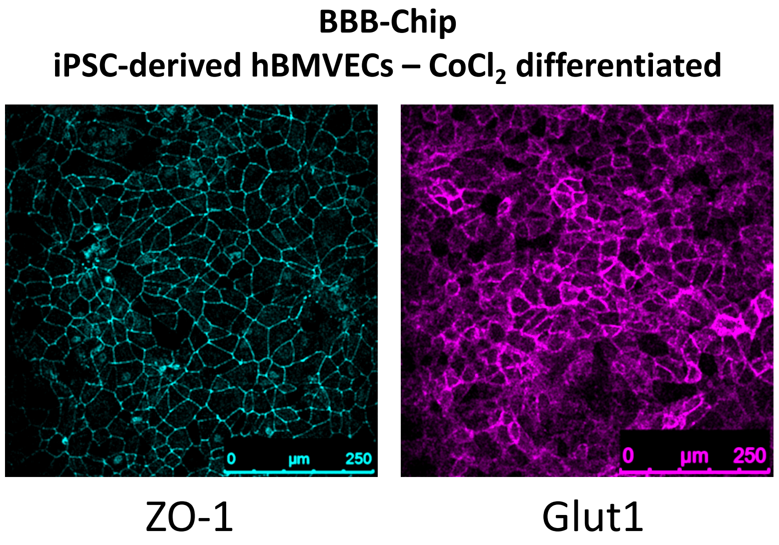


**Supplementary Fig. S9**

Immunofluorescence micrographs showing the distribution of tight junction protein, ZO-1, (left) and glucose transfer protein, Glut1, (right) on the surface of the endothelium within BBB chips generated by using iPS-BMVECs differentiated in the presence of $CoCl_2$.

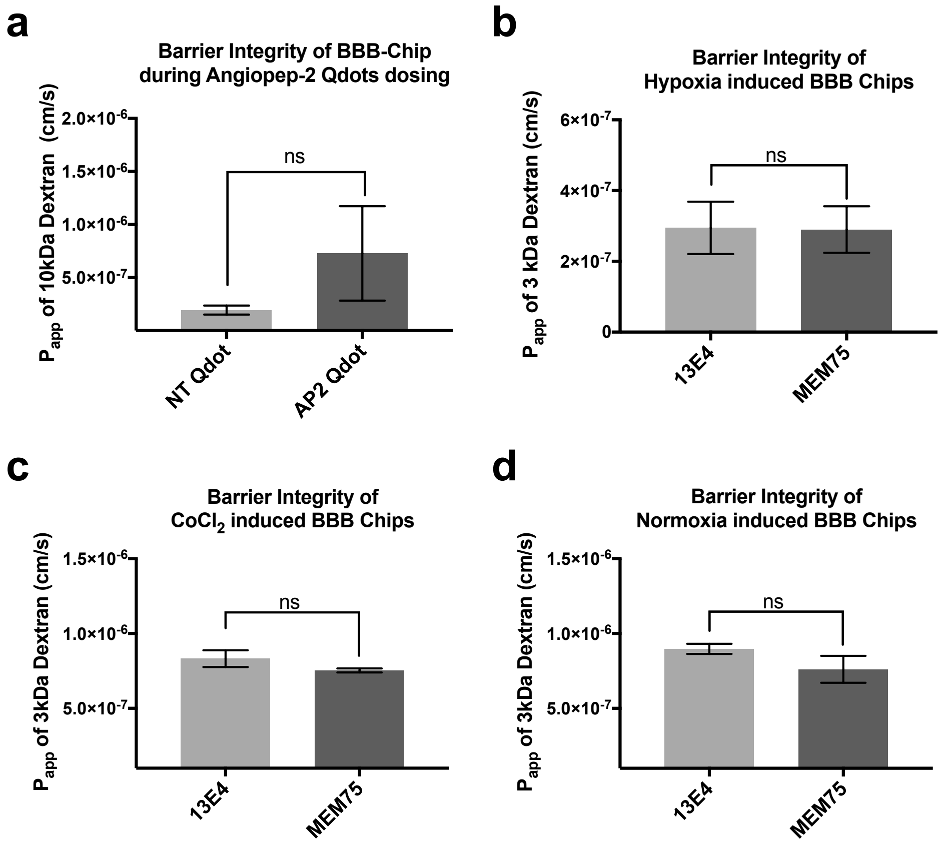


**Supplementary Fig. S10**

No statistically significant differences (ns) in barrier integrity were detected in the BBB Chips in experiments measuring transcytosis of Angiopep-2 **(a)** or anti-TfR antibodies (MEM75 and 13E4) in iPS-BMVECs induced by hypoxia **(b)**, $CoCl_2$ **(c)**, or normoxia **(d)** as monitored by measuring the permeability of 3 or 10 kDa dextran tracers.

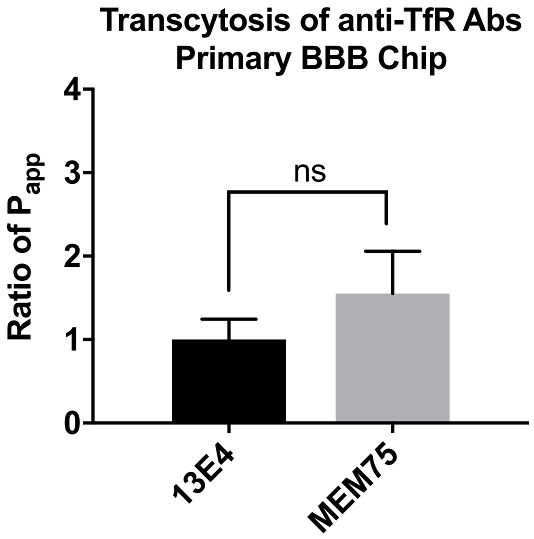


**Supplementary Fig. S11**

Transcytosis of anti-TfR antibodies, MEM75 and 13E4, measured by quantifying their relative apparent permeability (Ratio of $P_{app}$) in the primary BBB Chip, demonstrating that there was no significant difference (N.S.) between the transcytosis abilities of the two anti-TfR antibodies when primary human brain endothelial cells were used instead of iPS-BMVECs in the BBB Chip.

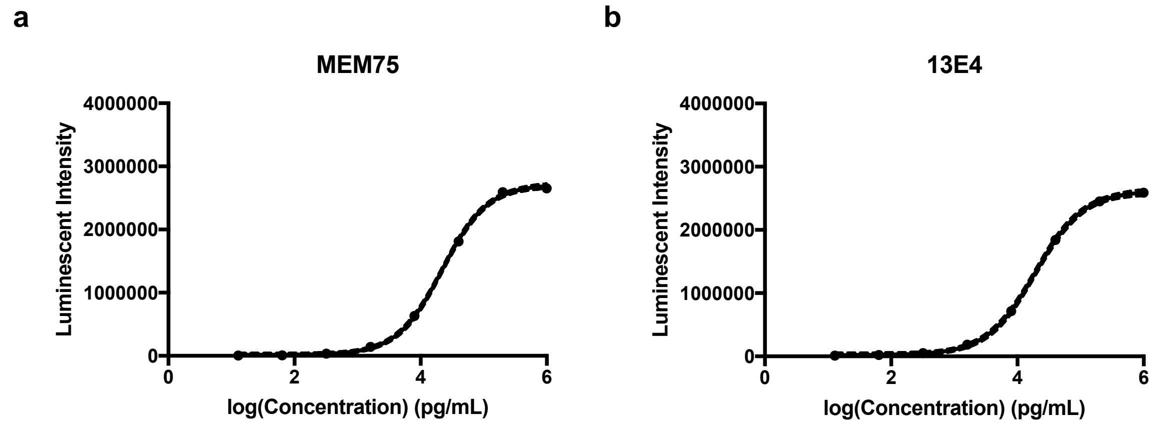


**Supplementary Fig. S12**

Standard curves for the binding of **(a)** MEM75 and **(b)** 13E4 antibodies to iPS-BMVECs that were used in the ELISA experiment.
